# The Genetic Basis of Chloride Exclusion in Grapevines

**DOI:** 10.1101/2025.02.28.640828

**Authors:** Sadikshya Sharma, Noe Cochetel, Jose Munoz, Hollywood Banayad, Yaniv Lupo, Veronica Nunez, Ana Gaspar, Christopher Chen, Krishna Bhattarai, Dario Cantu, Luis Diaz-Garcia

## Abstract

Mediterranean regions are among the most important areas for global grape production, characterized by dry climates and frequent challenges associated with soil salinity. In these environments, chloride toxicity is a major factor limiting vine growth and fruit quality. Despite the critical role of chloride exclusion in salinity tolerance, the genetic mechanisms underlying this trait remain poorly understood. In this study, we leveraged one of the largest *Vitis* germplasm collections, comprising 335 accessions from 18 wild and cultivated *Vitis* species, to characterize natural variation in chloride exclusion. This diverse panel, which includes accessions from the southwestern United States and Mexico, captures a broad range of evolutionary adaptations to abiotic stress, providing an unprecedented opportunity to investigate the genetic basis of salinity tolerance. Using genome-wide association (GWA) and quantitative trait loci (QTL) mapping, we identified a major QTL on chromosome 8 containing candidate genes encoding cation/H⁺ exchangers (CHX), which are involved in ion transport and homeostasis. To validate these findings, we analyzed a mapping population derived from *V. acerifolia* longii 9018 and the commercial rootstock GRN3, confirming the chromosome 8 locus as a major determinant of chloride exclusion. Structural variant analysis revealed key non-synonymous substitutions within CHX genes, suggesting potential functional roles in salinity tolerance. Additionally, we discovered a novel QTL on chromosome 19 enriched with G-type lectin S-receptor-like serine/threonine-protein kinases, known regulators of stress signaling. By integrating extensive phenotypic and genomic data across a diverse *Vitis* collection, this study provides novel insights into the genetic architecture of chloride exclusion and identifies valuable candidate genes for breeding salt-tolerant rootstocks.

## Introduction

Soil salinization is a major abiotic stress affecting agriculture worldwide. Approximately 1.4 billion hectares of land worldwide are affected by salinity, with an additional 1 billion hectares at risk (FAO, 2024). The primary contributors to soil salinization include inadequate irrigation practices, poor-quality irrigation water, overexploitation of water resources, and climate change-induced factors such as rising temperatures and altered precipitation patterns (Olesen et al., 2011; Ollat et al., 2016). Grape production is particularly relevant in semi-arid or Mediterranean-type climates, where limited rainfall and higher evaporation rates can exacerbate salt accumulation, making vines particularly vulnerable. Soil salinity not only reduces grape yield but also impacts fruit quality (Dobrei et al., 2016; Walker et al., 2003). Excessive sodium (Na^+^) and chloride (Cl^−^) accumulation in plant tissues can induce ion toxicity symptoms such as leaf necrosis and reduced photosynthesis, leading to stunted vegetative growth, leaf burn, and defoliation (Baby et al., 2016; Walker et al., 2010). Studies have shown that grapevines are moderately sensitive to salinity, with growth reductions starting at soil electrical conductivity levels of 1.5 dS/m, and yield losses of approximately 9.6% for each additional dS/m increase (Grieve et al., 2012). Beyond yield reductions, high soil salinity disrupts the balance of sugars, acids, and flavor compounds in grapes, ultimately affecting the resulting wine’s enological properties and quality (Dobrei et al., 2016; Walker et al., 2003). As climate change intensifies, the frequencies of hot and dry periods are expected to rise, exacerbating soil salinization and further threatening grape production (Hannah et al., 2013; Phogat et al., 2020; Walker et al., 2003; Munns and Tester 2008; Baby et al., 2016).

Grapevines employ multiple strategies to tolerate salt stress, including shoot ion exclusion (Henderson et al., 2014; Heinitz et al., 2020), vacuolar sequestration of intracellular ions (Wu et al., 2019), reactive oxygen species (ROS) signaling and detoxification (Zhang et al., 2016; Tanveer et al., 2020), and osmotic adjustment through accumulation of organic osmolytes (Sharma et al., 2019; Van Zelm et al., 2020). Salt exclusion, a key mechanism, enables plants to limit the uptake of Na^+^ and Cl^−^ from the soil and/or restrict their translocation from roots to shoots via the xylem (Wu et al., 2019). While the genetic and physiological mechanisms governing Na^+^ exclusion are relatively well understood (da Silva et al., 2024; Henderson et al., 2018; Mohammadkhani et al., 2016; Wu et al., 2020), Cl^−^ exclusion remains less characterized. Despite both Na^+^ and Cl^−^ contributing to salinity stress, grapevine rootstocks have traditionally been classified as tolerant or susceptible based on their ability to restrict Cl^−^ uptake, with Na^+^ exclusion considered a secondary factor (Heinitz et al., 2020; Walker et al., 1994a; Walker et al., 2003). This distinction is particularly significant in California, one of the world’s largest grape-producing regions, where Cl^−^ severely impacts many major viticultural areas, making chloride exclusion a key trait for the development of salt-tolerant rootstocks (Fort et al., 2013; Fort et al., 2015; Heinitz et al., 2020; Sharma et al., 2024)

Wild *Vitis* species exhibit greater variability in Cl^−^ exclusion than most commercially available rootstocks, highlighting their potential as a genetic resource for improving salt tolerance. A comprehensive survey of 325 accessions from 14 *Vitis* species collected across the southwestern United States and Mexico revealed substantial variation in Cl^−^ exclusion, both between and within species (Heinitz et al., 2020). Some accessions exhibited strong chloride exclusion —accumulating significantly less Cl⁻ than 140 Ruggeri, a widely used commercial rootstock for high-salinity conditions (Sharma et al., 2024)—while others showed weak exclusion ability, underscoring considerable intraspecific variability. Certain species, such as *V. acerifolia*, consistently restricted Cl^−^ uptake, while others displayed a broad range of chloride accumulation, suggesting that multiple genetic mechanisms regulate this trait (Heinitz et al., 2020; Sharma et al., 2024). Building on these findings, Cochetel et al. (2023) identified a genomic region on chromosome 8 (position 13,598,495 bp) associated with chloride exclusion, based on the Heinitz et al. (2020) phenotypic survey. They discovered a predicted protein homologous to the *Arabidopsis thaliana* cation/H⁺ exchanger AtCHX20 (*AT3G53720*) within this region. The putative function of this gene was further supported by the presence of two key InterPro domains: the Cation/H⁺ exchanger (IPR006153) and the Sodium/solute symporter superfamily (IPR038770). Moreover, the gene upstream of the significant SNP was also found to be homologous to *AtCHX20*, reinforcing its potential role in Cl^−^ exclusion. To date, no other grapevine candidate genes, major or minor, have been identified for Cl^−^ exclusion.

In this study, we explored the extensive genetic diversity of 335 accessions spanning 18 *Vitis* species and uncovered remarkable variation in Cl⁻ exclusion both within and between species. This broad survey underscores the rich genetic potential within wild and cultivated *Vitis* germplasm for enhancing salt tolerance. Building on these findings, our genome-wide association study (GWAS) and linkage mapping not only confirmed the previously reported QTL on chromosome 8 (Cochetel et al., 2023)—a region containing cation/H⁺ exchanger (CHX) genes—but also identified an orthologous locus in *V. acerifolia* that reinforces the importance of these candidate genes. In addition, we discovered a novel QTL on chromosome 19 enriched with *G*-type lectin S-receptor-like serine/threonine-protein kinase genes known to play significant roles in salt stress adaptation. Collectively, these results advance our understanding of the genetic underpinnings of chloride exclusion, highlighting the power of exploiting natural *Vitis* diversity to develop improved, salt-tolerant rootstocks and cultivars.

## Results

### Interspecies and intraspecies variation for chloride exclusion

We conducted 4 independent Cl^−^ trials, each evaluating between 61 and 141 accessions and 4 replicates per accession, over two years to assess 335 *Vitis* accessions spanning 18 species. In total, 1340 vines were examined. Chloride exclusion was measured as the amount of Cl^−^ in the leaves after 21 days of growth under 50 mM NaCl. The distribution of Cl⁻ exclusion was continuous and skewed toward lower chloride content (Figure 1A), with high variability observed both between and within species. Best linear unbiased predictors (BLUPs) ranged from 5.07 (OK14-006, *V. acerifolia*) to 197.45 mg/L (Vru 110, *V. rupestris*), with a mean of 69.45 mg/L. In addition, notable interspecific differences were observed (Figure 1B). Among those with the lowest mean values, *V. shuttleworthii* exhibited the lowest chloride content (23.0 mg/L), although this estimate is based on a single sample. Similarly, *V.* x *doaniana* (mean = 39.6 mg/L, n = 8) and *V. acerifolia* (mean = 46.4 mg/L, n = 17) also displayed low mean chloride levels. In contrast, species with higher chloride contents included *V. labrusca* (mean = 131.0 mg/L, n = 2), *V. rupestris* (mean = 126.0 mg/L, n = 10), and *V. californica* (mean = 88.6 mg/L, n = 6). The largest intraspecific variation was found in *V. labrusca* (SD = 72.1, n = 2), *V. rupestris* (SD = 48.8, n = 10), and *V. champinii* (SD = 36.7, n = 17), whereas *V. candicans* (SD = 11.8, n = 25), *V. aestivalis* (SD = 13.9, n = 4), and *V. vulpina* (SD = 21.7, n = 5) had the lowest. The number of accessions sampled per species did not correlate with the standard deviation (r = 0.05, p = 0.8333).

**Figure 1.**
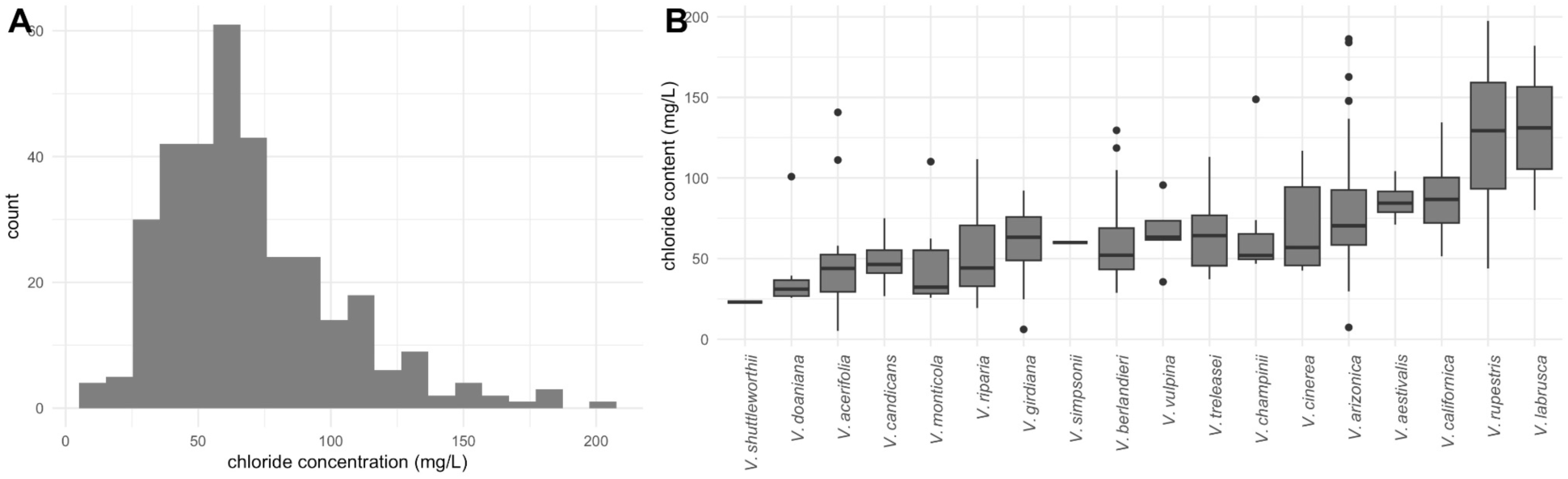
Variability in leaf Cl^−^ content across 335 accessions and 18 *Vitis* species. **(A)** Histogram of the average leaf Cl^−^ content using BLUPs for all the accessions. **(B)** Leaf Cl^−^ content across 18 *Vitis* species.

### Phylogenetic and population structure analysis

We combined whole-genome sequence data from two previous studies, which included 313 of the accessions used in our chloride screening (Cochetel et al., 2023; Morales-Cruz et al., 2021; Morales-Cruz et al., 2023). We performed a new SNP-calling analysis using the haplotype 1 genome of *V. giridiana* SC2 (Cochetel et al., 2023), yielding 2,336,705 markers (an average of 122,984 per chromosome) with a minor allele frequency (MAF) above 0.1.

We examined population structure and phylogeny to infer the demographic history of *Vitis* and its impact on chloride exclusion (Figure 2A). Genetic structure analysis revealed nine distinct ancestral genetic pools, which were differentially present among the individuals examined (Figure 2B). Based on this, several species, such as *V. berlandieri, V. candicans, V. californica,* and *V. rupestris*, formed well-defined groups with little to no admixture. Interestingly, even within these uniform groups, chloride exclusion varied by more than 4-fold for some species (e.g., *V. rupestris, V. berlandieri*), likely due to adaptive introgressions through hybridization or genetic differentiation driven by local adaptation. Other species, such as *V. vulpina, V. labrusca,* and *V. treleasei*, exhibited more mixed ancestry. The Neighbor-Joining tree (Figure 2C) further illustrated these relationships.

**Figure 2.**
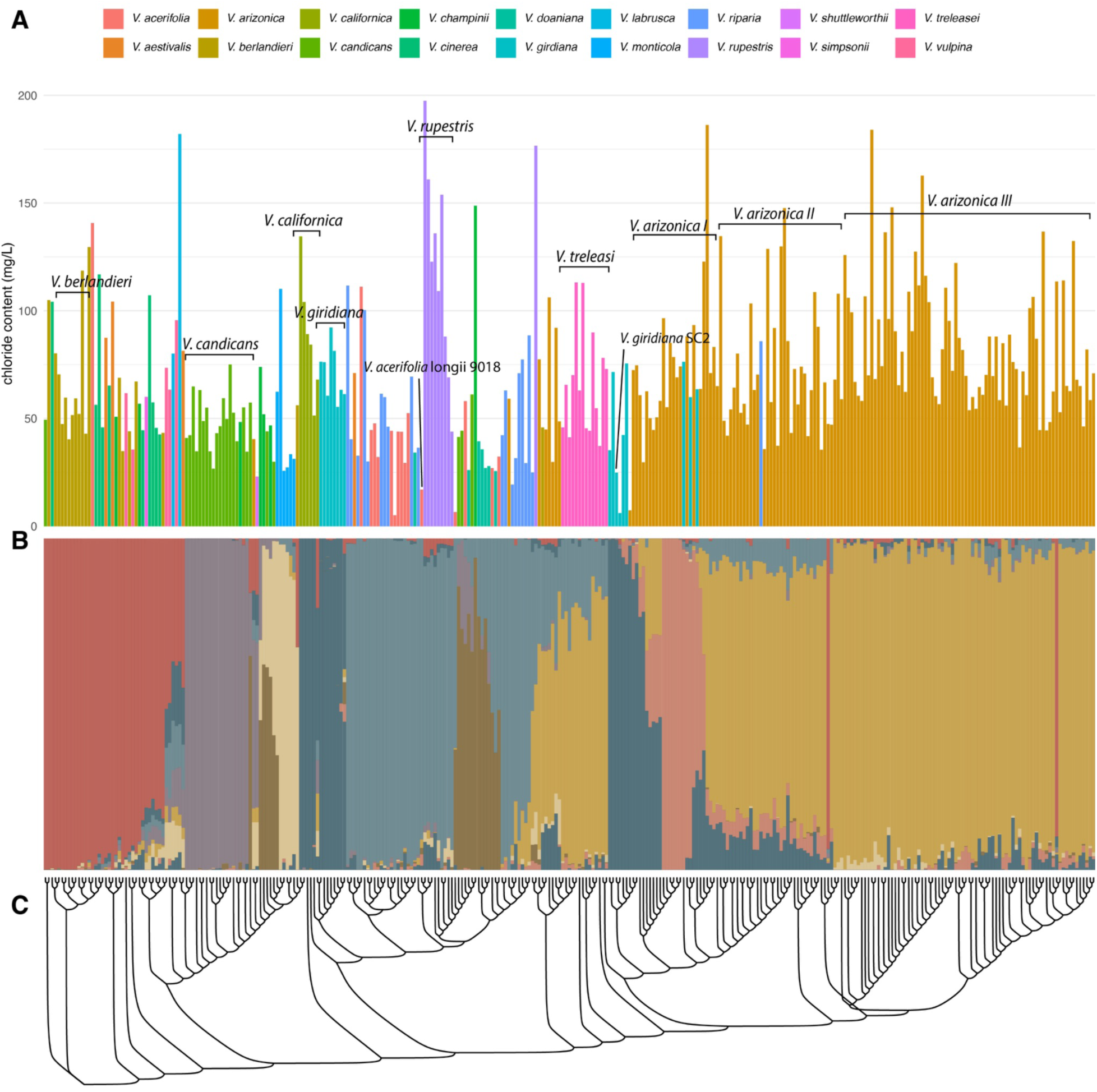
Population structure and phylogenetic relationships among 313 Vitis accessions with varying chloride exclusion. **(A)** Chloride content in each accession, arranged in the same order as the phylogeny in panel (C). Bars are colored according to species. **(B)** Structure analysis showing ancestry proportions for each accession (vertical bars), with colors representing different genetic clusters (unrelated to the species colors in panel A). Accessions are arranged as in panel (C). **(C)** Neighbor-Joining phylogenetic tree.

For 256 of the accessions, we compiled passport data providing the geographical coordinates of their sampling locations. We then used these coordinates to examine whether geographic separation—particularly within species—correlates with chloride exclusion (Figure 3A). Our hypothesis was that populations in closer proximity would encounter similar soil and climatic conditions, potentially leading to shared adaptive responses. However, we found diverse patterns in geographical distribution and Cl^−^ exclusion capabilties. For instance, most *V. treleasei* accessions came from relatively confined locations yet displayed considerable variability in chloride exclusion. Conversely, species such as *V. arizonica* and *V. riparia* spanned broad geographic ranges and covered nearly the entire spectrum of chloride exclusion found in this study. These findings suggest that geographic proximity alone does not fully explain the phenotypic diversity in chloride exclusion across *Vitis* species. We acknowledge that this approach has certain limitations. Soil conditions can vary substantially over short distances, which might explain large phenotypic responses across accessions relatively closer to each other.

**Figure 3.**
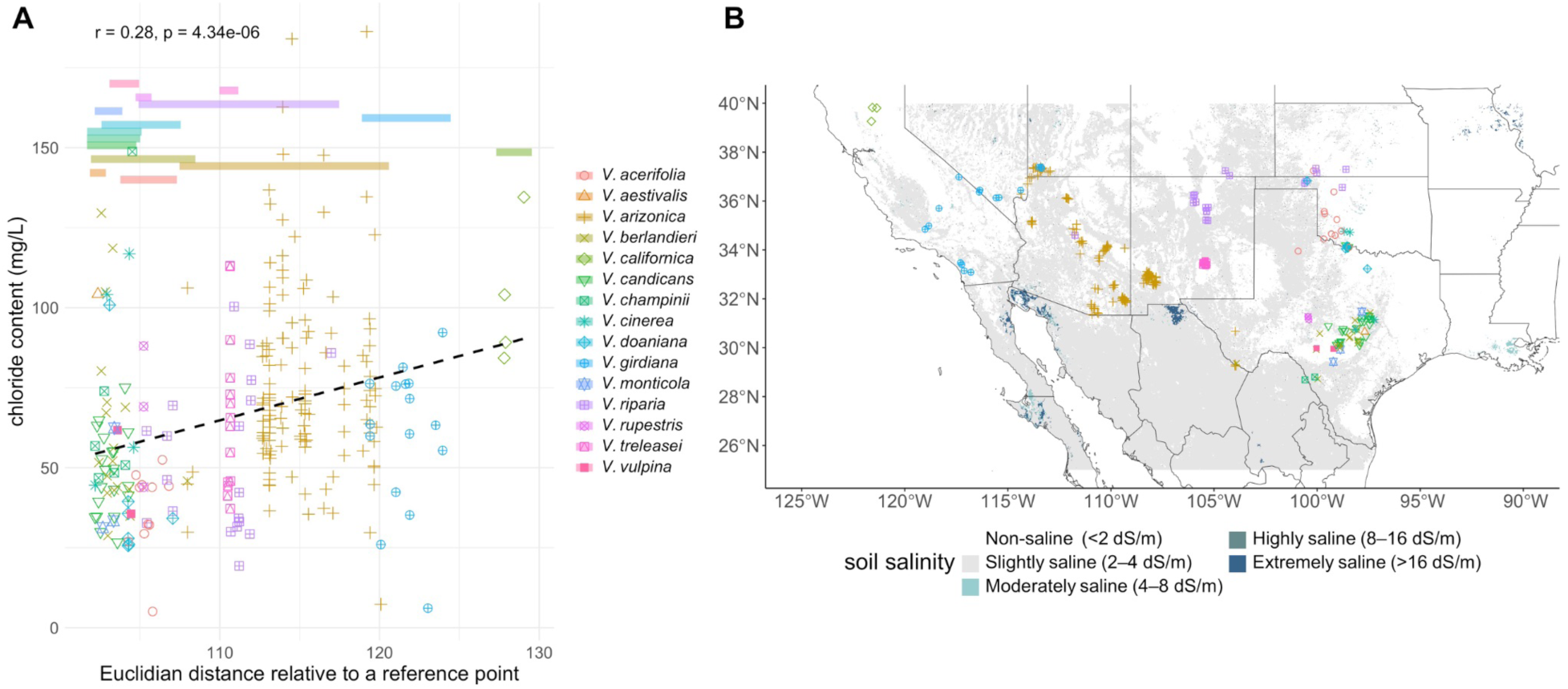
Geographic distribution, soil salinity, and chloride exclusion variation among 256 *Vitis* accessions. **(A)** Relationship between chloride content and Euclidean distance from a reference point (0,0), based on latitude and longitude. Each point represents an accession, colored and shaped according to its species. The dashed line indicates the correlation between geographic distance and chloride accumulation (r = 0.28, p = 4.34e-06). Horizontal bars at the top represent the geographical range of each species; their position along the y-axis is arbitrary and does not indicate specific chloride values. **(B)** Spatial distribution of accessions across the southwestern United States and northern Mexico, with point colors matching species assignments from panel A. Background shading represents soil salinity levels, classified as non-saline (<2 dS/m), slightly saline (2–4 dS/m), moderately saline (4–8 dS/m), highly saline (8–16 dS/m), and extremely saline (>16 dS/m).

To consider soil properties not accounted for by geography, we compiled soil salinity data from the global soil salinity map (Ivushkin et al., 2019), which was generated using a random forest classifier trained on soil property maps, thermal infrared imagery, and electrical conductivity (ECe) point data from the WoSIS database (Batjes et al., 2017). Rather than directly mapping Na⁺ or Cl⁻ concentrations, this dataset relies on ECe as an indicator of total dissolved salts, including sodium, chloride, and other ions. Soil salinity is classified into discrete categories: non-saline (<2 dS/m), slightly saline (2–4 dS/m), moderately saline (4–8 dS/m), highly saline (8–16 dS/m), and extremely saline (>16 dS/m). Among the *Vitis* accessions with available geographic sampling data, 114 were collected from sites classified as slightly saline (2–4 dS/m), while 142 were from non-saline locations (<2 dS/m; Figure 3B). We found non-statistical differences between leaf Cl^−^ content and soil salinity class (F = 0.9513, p = 0.3303). Several explanations can be made, including that this dataset does not truly capture the salinity conditions at the sampling locations of the accessions in this study, or that there are other forces driving chloride exclusion, such as climate. We then tested associations between climatic variables compiled with the “*EnvRtype”* package (Costa-Neto et al., 2021), and chloride exclusion levels. Specifically, mean temperature of the driest quarter (F = 24.76, p = 1.21e^−06^), max temperature of the warmest month (F = 11.79, p = 0.000696), and mean temperature of the warmest quarter (F = 14.64, p = 0.000164) were all strongly associated with chloride exclusion, suggesting that heat stress during drier periods may play a role in chloride adaptation. Similarly, precipitation patterns were also relevant, with precipitation of the coldest quarter (F = 11.44, p = 0.000835), precipitation of the driest quarter (F = 11.15, p = 0.000965), precipitation seasonality (F = 11.51, p = 0.000802), and precipitation of the driest month (F = 14.08, p = 0.000218) all showing significant associations with chloride exclusion levels (data not shown).

We note that our inferences about *Vitis* chloride exclusion phenotypes are shaped by the genetic and geographic sampling included in this study, which could introduce bias—particularly for under sampled species. Additionally, given the interspecific compatibility of *Viti*s species and their ability to produce fertile hybrids, some degree of misclassification is possible. Lastly, we emphasize that conducting a comprehensive phylogenetic analysis of *Vitis* was beyond the scope of this study.

### GWAS revealed loci and candidate genes associated with chloride exclusion

GWAS identified a single variant on chromosome 19 (position 4,769,399 bp) significantly associated with chloride exclusion (beta = 24.75 mg/L; Figure 4A), representing a novel QTL. At a more relaxed threshold −log10(p) = 6 (instead of −log10(p) = 7.58, corresponding to the FDR-adjusted threshold), we detected three additional associations on chromosomes 2 (8,523,304 bp, beta = 22.72 mg/L), 7 (21,049,011, beta = –18.89 mg/L), and 8 (13,602,850, beta = –29.78 mg/L). The chromosome 2 and 7 loci have not been previously reported, whereas the chromosome 8 locus coincides with a known QTL at the same region (13,598,495 bp; Cochetel et al., 2023). To account for population structure, we incorporated both a kinship matrix (from the full marker set) and 10 principal components (PCs). A Q–Q plot confirmed that these corrections mitigated population stratification (Figure 4B).

**Figure 4.**
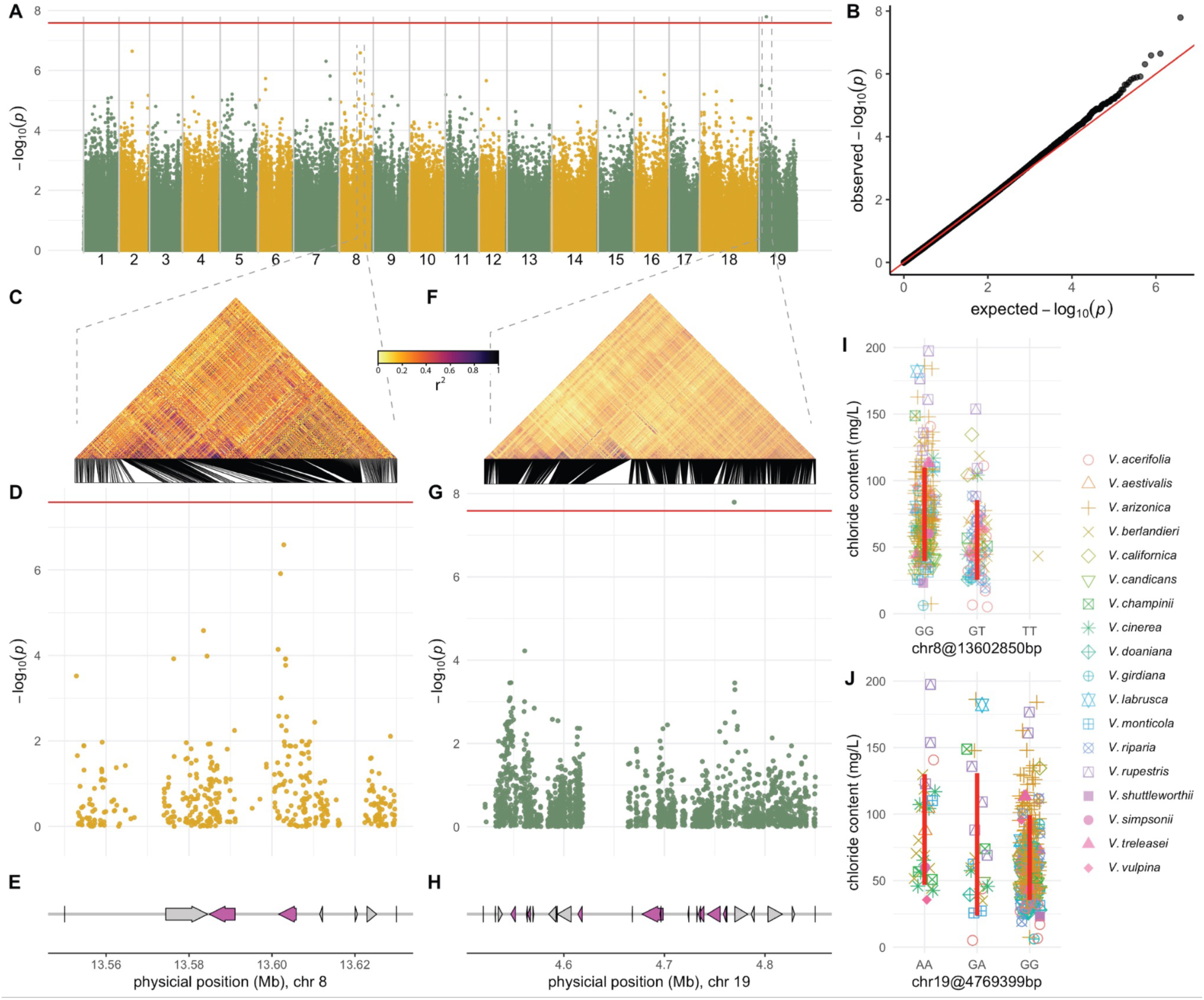
Genome-wide association of chloride exclusion in grapevine. **(A)** Manhattan plot displaying the genome-wide distribution of SNP associations (1,916,527 markers) with chloride exclusion across the grapevine genome. The horizontal red line denotes the significance threshold (FDR-adjusted p-value = 0.05), highlighting one significant quantitative trait loci. **(B)** Quantile-quantile plot assessing the observed versus expected −log₁₀(p) values under the null hypothesis, indicating deviations due to associations. **(C)** Linkage disequilibrium (LD) heatmap (r²) in the significant region of chromosome 8. **(D)** Local Manhattan plot for chromosome 8, zooming in on the significant region identified in (A). **(E)** Gene content in the significant chromosome 8 region, with genes represented as arrows. Genes of interest *(Cation/H⁺ Exchanger 3)* are highlighted in purple. LD heatmap (r²) for the significant region on chromosome 19, analogous to panel (C). **(G)** Local Manhattan plot for chromosome 19, highlighting significant associations. **(H)** Gene content in the significant chromosome 19 region, similar to panel (E), where genes are represented as arrows. Multiple copies of the G-type lectin S-receptor-like serine/threonine-protein kinase gene are highlighted in purple. **(I, J)** Effect plots displaying chloride content (mg/L) across different genotypes for the SNPs at chromosome 8, position 13.6 Mb and chromosome 19, position 4.79 Mb. Vertical red lines correspond to 2 standard deviations center in the mean by genotype.

Examination of the chromosome 8 QTL revealed a cluster of linked SNPs with rapid LD decay (Figure 4C), especially among those with the highest −log10(p) values (Figure 3D). As reported by Cochetel et al. (2024), this region contains two genes (*g144880*, 13,584,884–13,591,103; and *g144890*, 13,601,610–13,605,816) encoding predicted homologs of the Arabidopsis cation/H^+^ exchanger *AtCHX20* (*AT3G53720*; Figure 4E). A similar LD pattern was observed around the novel QTL on chromosome 19 at 4,769,399 bp (Figure 4F). The upstream region (Figure 4G) contains five copies (*g307480* at 4,562,881–4,563,707; *g307550* at 4,667,975–4,668,211; *g307570* at 4,695,533–4,696,547; *g307590* at 4,732,332–4,732,826; *g307640* at 4,761,909–4,762,151) of a gene predicted to encode a G-type lectin S-receptor-like serine/threonine-protein kinase homologous to *At4g27290* (Figure 4H). Four additional genes annotated as G-type lectin S-receptor-like kinases (*g307500, g307610, g307620, g307630*) were also identified in this region. This region is promising, as G-type lectin S-receptor-like serine/threonine-protein kinases have been identified as positive regulators of salt stress tolerance in multiple crops (Sun et al., 2013; Joshi et al., 2010; Sun et al., 2018). A full list of genes in the QTL intervals of chromosomes 2, 7, 8 and 19 is provided in Table S3.

### QTL mapping identifies two cation/H⁺ exchanger genes

To further investigate the genetic control of chloride exclusion in grapevines, we generated a mapping population derived from the chloride-excluding accession *V. acerifolia* longii 9018 and the commercial rootstock GRN3. longii 9018 has exceptional chloride exclusion capabilities (chloride content: 16.97 mg/L, based on the GWAS screening, Figure 2A). It has previously been reported as a chloride excluder (Sharma et al., 2024; Heinitz et al., 2020), making it a strong candidate for this study. GRN3 is a nematode-resistant rootstock with moderate salinity tolerance (Andrew Walker, personal communication) and has *V. champinii, V. rufotomentosa,* and *V. riparia* in its parentage.

The longii 9018 × GRN3 mapping population (n=162), referred to here as 18113, was genotyped using genotyping-by-sequencing, with variant calling based on the genome of *V. acerifolia* longii 9018 (Cochetel et al., 2023). Two genetic maps were constructed using a pseudo-test cross linkage mapping strategy (Grattapaglia and Sederoff, 1994). The parental map for longii 9018, which contained polymorphic markers (ABxAA or lmxll), included 2,611 markers across 740 bins (unique genetic positions). The parental map for GRN3 contained 4,842 markers across 784 bins. Physical and genetic distances showed strong agreement for both parental maps (Figure 5A).

**Figure 5.**
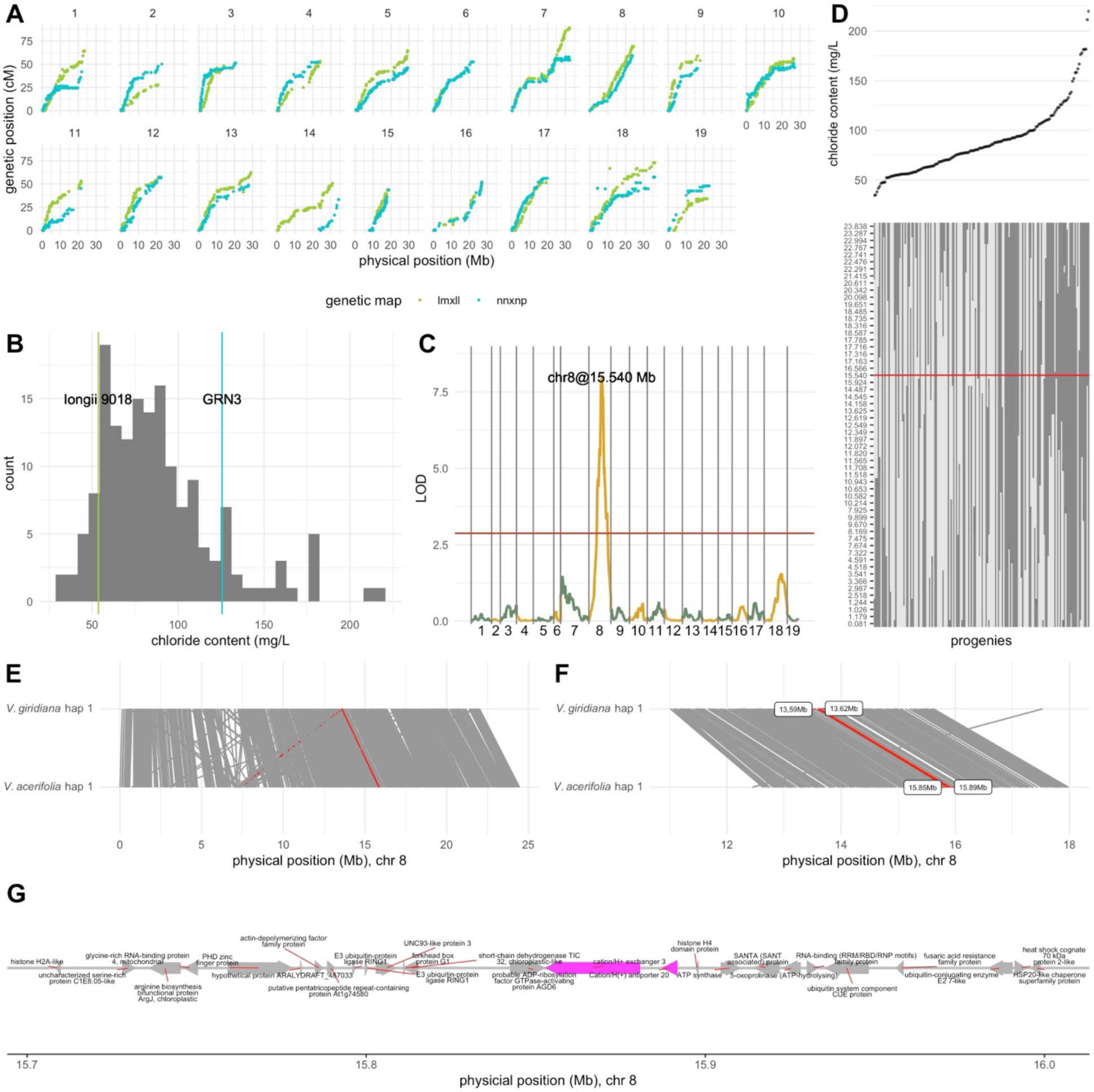
QTL mapping of chloride exclusion in the longii 9018 × GRN3 mapping population. (A) Relationship between genetic and physical positions for each chromosome in the parental maps (lm × ll in green, nn × np in blue). **(B)** Distribution of chloride content in the mapping population, estimated using BLUPs. The parental values are indicated with vertical lines: longii 9018 (green) and GRN3 (blue). **(C)** QTL scan for chloride exclusion based on the parental lm × ll map. The logarithm of odds (LOD) scores along the genome are shown, with a peak detected on chromosome 8 at 15.540 Mb (chr8@15.540 Mb). The red horizontal line represents the significance threshold obtained from 1,000 permutations at the 95th percentile. **(D)** Segregation analysis of chromosome 8 genetic markers and their relationship with chloride exclusion. Progenies (columns) are sorted by increasing chloride content (top panel). The genotype matrix (bottom panel) represents genotypic segregation across chromosome 8, with dark gray indicating homozygous (ll) and light gray indicating heterozygous (lm) individuals. The red horizontal line corresponds to the major QTL peak from panel (C). **(E)** Synteny along chromosome 8 between *V. giridiana* (haplotype 1) and *V. acerifolia* (haplotype 1). **(F)** Zoomed-in view of the syntenic region identified through GWAS, which contains the two *cation/H^+^ exchanger 3* genes. **(G)** Gene content based on the *V. acerifolia* longii 9018 genome (haplotype 1; used for SNP calling) across the candidate region (15.45–15.65 Mb).

The 18113 mapping population was screened for chloride exclusion using the same method as above (Heinitz et al., 2020; Sharma et al., 2024). The screening was conducted in a semi-augmented design across 4 trials, with varying numbers of genotypes per trial (n = 21, 21, 55, and 143). Among the progeny, 87 were included in only one trial, while 73 were tested in two trials. longii 9018 was included in all trials, whereas GRN3 was included in three. Transgressive segregation was observed, with chloride content ranging from 34.46 to 219.96 mg/L and a mean of 88.69 mg/L (Figure 5B). The chloride contents were 53.62 mg/L for longii 9018 and 125.61 mg/L for GRN3. The difference in chloride accumulation for longii 9018 between this study and the GWAS trial is not unexpected, as variability is observed across trials, highlighting the importance of including controls.

We used the scanone function from the R package qtl (Broman et al., 2003) to identify QTLs in each parental map separately. A major QTL was detected on chromosome 8 at 41.16 cM, with a LOD score of 8.01, explaining 20.6% of the variance (Figure 5C). This QTL was found on the parental map with markers segregating for longii 9018 (ABxAA or lmxll). Further examination of the segregants and their phenotypes at the QTL showed lower chloride content in heterozygous genotypes (Figure 5D, lighter gray) and higher chloride content in homozygotes (darker gray). The QTL marker was located at 15,540,293 bp in the longii 9018 haplotype 1 genome. Just downstream (within the 1.5-LOD support interval), between 15,853,017 and 15,891,666 bp, two contiguous genes were identified with predicted annotations: *Cation/H^+^ Exchanger 3* (*g154540*: 15,853,017–15,880,628) and *Cation/H(+) Antiporter 20* (*g154550*: 15,887,484–15,891,666). Using minimap2 (Li, 2018), we found that this region (chromosome 8, 15,853,017–15,891,666) is orthologous to the longii 9018 region in 13.60Mb containing the two *Cation/H^+^ exchangers* identified through GWAS (Figure 4). No QTLs, or any sign of significance, were found using GRN3 parental map, which indicates that this chloride exclusion locus is inherited from the *V. acerifolia* longii 9018 parent.

### Genetic variants located in the *cation/H⁺ AtCHX20 exchanger* gene

Considering the agreement between GWAS and QTL mapping, we further inspected the QTL region 13.56–13.62 Mb on chromosome 8 using an unfiltered (MAF > 0.05) version of the VCF for this region. Using GWAS, 5,216 markers were tested for association with chloride content. Kinship and 10 PCs derived from the genome-wide marker dataset were included in the GWAS model. The most significant markers were found in the second copy (*g144890*, 13,601,610–13,605,816) of the predicted homologs of the Arabidopsis cation/H⁺ exchanger *AtCHX20* (Figure 6A). Four SNPs (13,602,044; 13,602,071; 13,602,850; 13,602,955) within this gene had significantly higher −log₁₀(p) values (>5.5) compared to other surrounding markers. Minor allele frequencies for these SNPs ranged from 0.062 to 1.131. The *g144890* gene contains 4 exons, with the first two associated with the Sodium/Hydrogen Exchanger Transmembrane Domain (Pfam entry: PF00999) and the last two with the Plant Cation/H⁺ Antiporter Domain (Pfam entry: PF23256). All 4 SNPs were in exons 3 and 4. Among these SNPs, three resulted in nonsynonymous amino acid changes: D to E (13,602,044, G/C), M to L (13,602,850, C/A), and I to F (13,602,955, A/T). The 13,602,071 (G/A, R/R) SNP resulted in a synonymous mutation with no amino acid change. Marker effects (Figure 6B) for the three nonsynonymous changes were equal to or more pronounced than those observed in the GWAS with the genome-wide marker set (Figure 4I).

**Figure 6.**
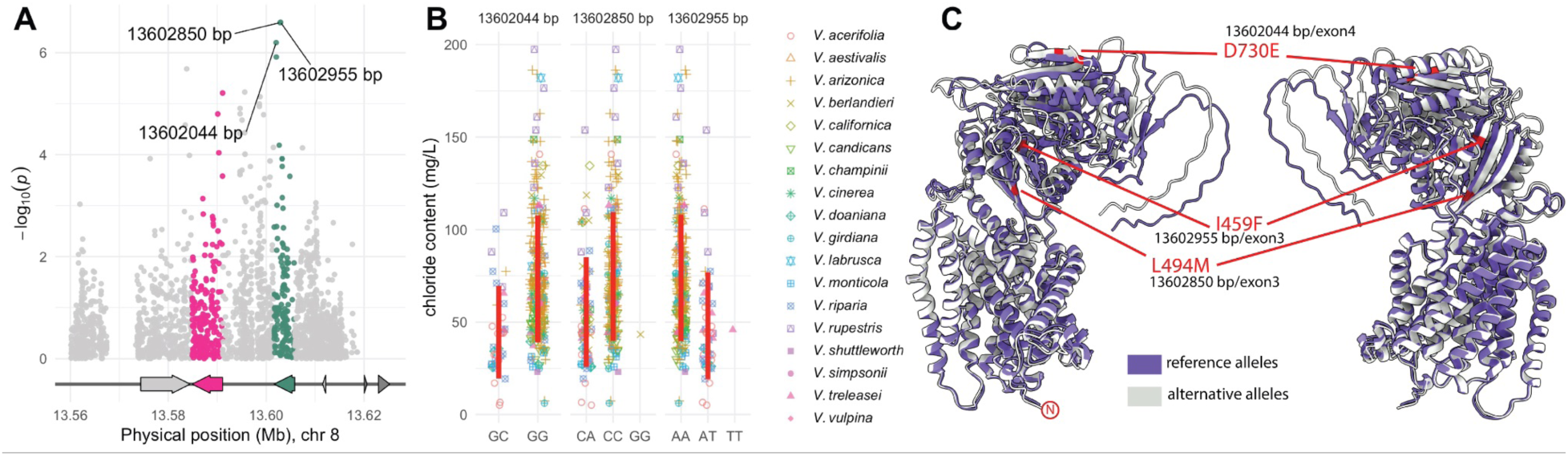
Identification of exonic variants in the cation/H⁺ *AtCHX20* exchanger gene associated with chloride exclusion. **(A)** Manhattan plot displaying the association of SNPs on chromosome 8 (13.56–13.62 Mb) with chloride content. The most significant markers are located in the second copy (*g144890*, green arrow) of a predicted cation/H⁺ exchanger homolog, with three SNPs (13,602,044 bp; 13,602,850 bp; 13,602,955 bp) showing high −log₁₀(p) values. The *g144880* gene and markers within it are colored in pink. **(B)** Effect plots displaying chloride content (mg/L) across different genotypes for three nonsynonymous changes at chromosome 8 for SNPs 13,602,044, 13,602,850, and 13,602,955. Vertical red lines correspond to 2 standard deviations center in the mean by genotype. **(C)** Structural model of the predicted cation/H⁺ exchanger protein, comparing the reference allele conformation (purple) with the alternative allele conformation (gray). The three nonsynonymous mutations (D730E, L494M, and I459F) are highlighted in red, indicating their positions within the exon 3 and exon 4 regions of *g144890*.

To further investigate the structural and functional consequences of these nonsynonymous substitutions, we used AlphaFold (Jumper et al., 2021) to predict the protein structures of the reference allele (wild type for all three SNP positions) and the alternative allele (mutant for all three SNP positions). Visually, no major structural alterations were observed between the two predicted protein conformations (Figure 6C).

Although the overall structure remained unchanged, individual amino acid substitutions may still influence local interactions. The aspartate (D) to glutamate (E) change is conservative, as both residues are negatively charged. However, glutamate’s longer side chain may introduce minor steric effects that could influence interactions, particularly in tight or functionally critical regions. The methionine (M) to leucine (L) substitution is also conservative due to their shared hydrophobic nature, but the loss of methionine’s sulfur-containing side chain could affect redox sensitivity or metal-binding properties if relevant to the protein’s function. In contrast, the phenylalanine (F) to isoleucine (I) change is less conservative, as it removes an aromatic ring, potentially disrupting π-stacking interactions or ligand binding. This alteration could also reduce local stability in regions where aromatic interactions are structurally significant. While these changes appear relatively mild, their potential functional consequences and role in differential chloride affinity require further investigation.

The 4,560,000-4,900,000 bp region of chromosome 19 was also inspected using 21,789 unfiltered SNP variants. However, the most significant marker was located in an intergenic region (Figure S1), and no further candidate gene search analyses were pursued.

## Discussion

### The diversity in chloride exclusion and the role of hybridization and local adaptation

The extensive phenotypic variation in leaf chloride content across *Vitis* species underscores the complexity of chloride exclusion, arising from diverse genetic and physiological mechanisms. Additionally, environmental factors such as soil composition, climate variability, and biotic interactions further influence these processes, contributing to the species’ differential responses to salinity stress. Here, we detected a broader range of chloride exclusion than previously reported. Our findings corroborate earlier observations regarding species that excel (*V. acerifolia*, *V. × doaniana*) and those that perform poorly (*V. rupestris*, *V. labrusca*) (Fort et al., 2013; Gong et al., 2011; Heinitz et al., 2020; Sharma et al., 2024). Recognizing the large intraspecific diversity is crucial, especially since traits like salinity tolerance are often viewed as fixed, species-level attributes (Rahemi et al., 2022).

Interspecific compatibility in *Vitis*, coupled with overlapping habitats, actively promotes hybridization—a key driver of phenotypic and structural genomic variability (Callen et al., 2016; Morales-Cruz et al., 2021; Cantu et al., 2024). In southwestern North America, multiple wild *Vitis* species frequently co-occur in canyon or riparian corridors, facilitating spontaneous introgression and admixture. Recent analyses of haplotype-resolved genome assemblies and a pangenome have revealed pervasive hemizygosity and structural variation—such as large inversions, translocations, and gene copy-number changes—across diverse *Vitis* lineages (Cochetel et al., 2023). This structural variation can directly impact essential phenotypic traits, including responses to soil salinity and other adaptation traits. These admixture patterns, while beyond the scope of this study, raise important questions about the degree to which physical barriers (e.g., desert basins, mountains) and local edaphic or climatic factors limit gene flow among overlapping wild populations. It also underscores cases wherein regionally confined species (e.g., *V. treleasei*) still display phenotypic breadth on par with widely distributed taxa like *V. arizonica*. Determining whether local gene flow among co-occurring species outweighs soil-driven or climate-driven selection requires further investigation.

### Two independent experiments point out cation/H+ exchanger 3 as a candidate gene for chloride exclusion in grapevines

Salinity tolerance in grapevines involves multiple genetic loci, regulating sodium and chloride uptake, transport, and tissue partitioning (Wu et al., 2019). Historically, research has focused more on sodium exclusion, given its critical role in major crops (Munns and Tester, 2008; Henderson et al., 2018). However, grapevines are often more susceptible to chloride toxicity, which manifests as marginal leaf burn, reduced photosynthesis, compromised fruit quality, and overall vine decline (Walker, 1994; Tregeagle et al., 2010). Consequently, the ability to exclude or compartmentalize chloride from shoot tissues is widely recognized as a key determinant of salinity tolerance in *Vitis* (Antcliff et al., 1983; Sykes, 1985; Fort et al., 2015; Heinitz et al., 2020). Despite significant progress in identifying sodium-exclusion mechanisms—most notably *VvHKT1;1*, which functions in retrieving sodium from xylem vessels into pericycle cells (Henderson et al., 2018)—the genetic basis of chloride exclusion has remained largely elusive. Several hypotheses have been proposed, including SLAH1– SLAH3-mediated chloride efflux at the root pericycle (Cubero-Font et al., 2016), direct chloride extrusion from roots (Abbaspour et al., 2013; Wu et al., 2019), and shoot-to-root recirculation of chloride (Godfrey et al., 2019; Qu et al., 2021). The relative complexity of these processes may explain why both single-locus (Antcliff et al., 1983) and polygenic (Sykes, 1985; Fort et al., 2015) models have been suggested for chloride exclusion in grapevines.

Recently, Cochetel et al. (2023) identified a QTL associated with chloride exclusion on chromosome 8, which colocalized with two genes encoding proteins homologous to the *A. thaliana* cation/H⁺ exchanger AtCHX20 (*AT3G53720*). Our study, using two independent mapping approaches—GWAS and QTL mapping—confirmed this locus with a dataset nearly twice the size of that used by Cochetel et al. (2023) and employing two different platinum-level genomes for variant calling (*V. girdiana* and *V. acerifolia*). AtCHX20 is a member of the cation/proton antiporter (CPA2) family in Arabidopsis, primarily involved in ion homeostasis and pH regulation within plant cells. It plays roles in guard cell movement, osmoregulation, and membrane trafficking (Padmanaban et al., 2007; Chanroj et al., 2012). Phylogenetic analyses suggest that *AtCHX20* and its homologs originated from cyanobacterial NhaS4-like genes and diversified in early land plants, leading to specialized functions in ion transport and cellular adaptation to environmental stress (Sze et al., 2004). In Arabidopsis, *AtCHX20* localizes to endomembranes, where it influences vesicular trafficking and intracellular pH balance—key processes that aid in plant adaptation to fluctuating environmental conditions. Other family members, such as *AtCAX3* and *AtCAX4*, have been shown to be upregulated under salt stress (Cheng et al., 2002), while the overexpression of rice CAX (*OsCHX11*) improved salt tolerance (Senadheera et al., 2009). In soybean, *GmCHX1* was also identified as a major gene conferring salt tolerance (Do et al., 2019).

The discovery that *Vitis* homologs of *AtCHX20* colocalize with a QTL for chloride exclusion suggests a potential role of cation/proton exchangers in chloride homeostasis, either through compartmentalization or by modulating ion fluxes that indirectly affect chloride transport. However, based on findings from Heinitz et al. (2020) and Cochetel et al. (2023), it is evident that this locus on chromosome 8 does not regulate chloride root compartmentalization. Chloride concentrations in root tissues showed no correlation with either leaf chloride concentrations or the genotype at this locus, reinforcing the idea that the mechanism of exclusion linked to this locus does not involve root sequestration.

The identification of what appears to be a large-effect QTL for chloride exclusion on chromosome 8 represents a big opportunity in rootstock breeding, as it provides an avenue for rapid allele introgression through marker-assisted selection (MAS). Historically, grape breeding has prioritized single-gene traits primarily in scion breeding, focusing on traits such as resistance to powdery mildew (Coleman et al., 2009; Pap et al., 2016), downy mildew (Bhattarai et al., 2021; Sapkota et al., 2023; Schwander et al., 2012), and Pierce’s disease (Krivanek et al., 2006; Riaz et al., 2006), as well as improvements in fruit quality (Chen et al., 2015; Wang et al., 2020; Reshef et al., 2022). Many of these traits have been introgressed from wild *Vitis* species into *V. vinifera* backgrounds (Morales-Cruz et al., 2023; Svyantek et al., 2020; Thomas et al., 2020). In contrast, candidate gene discovery for rootstock traits has been far more limited, with only a few notable exceptions, such as *XiR1*, which confers resistance to dagger nematode (Xu et al., 2008). The discovery of a genetic locus associated with chloride exclusion provides a critical tool for developing rootstocks with enhanced salinity tolerance, surpassing what is currently achievable with commercially available options like 140Ru. The ability to introgress this trait efficiently into elite rootstock germplasm through MAS could reduce the time and cost required to develop new, more resilient varieties. Additionally, understanding the functional mechanism of CHX-type transporters in chloride exclusion could inform transgenic or gene-editing approaches to further optimize rootstock performance under saline conditions.

### Novel QTLs for chloride exclusion

While other weak and strong associations were identified in our GWAS, the QTL on chromosome 19 stands out as particularly promising. This region contains nine gene copies predicted to encode G-type lectin S-receptor-like serine/threonine-protein kinases, which have been previously documented as key regulators of ion homeostasis and stress signaling in *Vitis* (Han & Li, 2024). These kinases play crucial roles in plant responses to environmental stressors, including salinity, by mediating signal transduction pathways that regulate ion transport, oxidative stress responses, and osmotic balance. Previous studies provide strong evidence for the involvement of receptor-like kinases (RLKs) in salt stress responses. For instance, the Ca²⁺-dependent protein kinase VaCPK21 in *V. amurensis* is significantly upregulated under salt stress, suggesting a role in stress adaptation (Dubrovina et al., 2016). More specifically, G-type lectin S-receptor-like kinases have been identified as positive regulators of salt tolerance across multiple species, including Arabidopsis (Sun et al., 2013), pea (Joshi et al., 2010), and alfalfa (Sun et al., 2018). In alfalfa, overexpression of *GsSRK-f* and *GsSRK-t* enhanced salt tolerance by modulating ion homeostasis, ROS scavenging, and osmotic regulation (Sun et al., 2018). Similarly, overexpression of *PsLecRLK* in tobacco improved Na⁺ compartmentalization and ROS detoxification, thereby increasing salinity tolerance (Vaid et al., 2015). In Arabidopsis, *AtLecRK2* and *AtLecRK-b2* are upregulated in response to salt and osmotic stress (He et al., 2004; Deng et al., 2009), while in rice, G-LecRLKs show increased expression under cold, drought, and salt stress (Vaid et al., 2012).

The identification of G-LecRLK in our study suggests that this gene family may contribute to grapevine adaptation to salt stress by modulating stress-responsive signaling cascades. Plants employ a combination of strategies to cope with salinity, including ion exclusion, osmotic regulation, and oxidative stress management (Xiong et al., 2002). The genes identified in this study—*CHX3* and *G-LecRLK*—may function in concert to enhance chloride exclusion and overall salt tolerance in grapevines. Alternatively, they may represent distinct adaptation mechanisms that vary across species or even among individual accessions, highlighting the complexity of salinity tolerance in *Vitis*. Further functional validation of these loci will be essential to unravel their precise roles in grapevine salt stress responses.

Future research should focus on validating the functional role of CHX homologs in *Vitis*, possibly through transcriptomic, physiological, and CRISPR-Cas9 knockout studies. Additionally, further work is needed to investigate the potential interplay between this chloride-exclusion mechanism and other abiotic stress responses, including drought and nutrient uptake efficiency. Given that chloride exclusion mechanisms in *Vitis* remain underexplored, the findings of this study set the stage for further advances in both fundamental and applied plant breeding research.

## Materials and methods

### Plant material

The accessions examined in this study are part of the University of California, Davis germplasm collection, which has been screened for disease resistance (Morales-Cruz et al., 2023), abiotic stress tolerance (Morales-Cruz et al., 2021), and genome composition (Cochetel et al., 2023). Over the years, the genomes of individuals in this collection have been increasingly characterized at multiple levels, ranging from the development of platinum-level reference genomes (Minio et al., 2022; Cochetel et al., 2023) to whole-genome sequencing (Morales-Cruz et al., 2021; Morales-Cruz et al., 2023) and the construction of a multi-species pangenome (Cochetel et al., 2023). In this study, we analyzed 335 accessions representing 18 *Vitis* species, primarily from the southwestern United States. The species included *V. acerifolia*, *V. aestivalis*, *V. arizonica*, *V. berlandieri*, *V. californica*, *V. candicans*, *V. champinii*, *V. cinerea*, *V. × doaniana*, *V. girdiana*, *V. monticola*, *V. riparia*, *V. rupestris*, *V. treleasei*, and *V. vulpina*. These accessions are clonally preserved at the University of California, Davis campus, in Davis, California, USA. Passport data and chloride content is available in Table S2.

### Experiment design

Green cuttings from the 335 accessions were collected during the spring over two growing seasons (2023 and 2024). Collections took place in the early morning (6:00–10:00 AM) to ensure tissue viability. To enhance rooting, cuttings were treated with 2% indole-3-butyric acid (IBA) before being placed in trays filled with pre-soaked perlite. The trays were maintained in a fog room (100 % relative humidity and 27 °C) for 14 days to promote root development and bud emergence. Once rooted, the cuttings were transplanted into 4-inch pots containing fritted clay and grown for two months before initiating the chloride stress treatment. Previous studies have established 50 mM NaCl as an appropriate concentration to induce salinity stress in potted grapevines using a similar screening method (Heinitz et al., 2020; Sharma et al., 2024). To ensure consistent salt exposure, each pot received 1 L of saltwater solution daily between 7:00 and 8:00 AM. Given the large number of accessions and replication levels, chloride exclusion screening was conducted across 4 independent trials using an augmented design. Four accessions—140 Ru, longii 9018, St. George, and 44-53 M—were included as controls in all trials, as they have been previously characterized as either chloride excluders or non-excluders. The number of accessions and key dates for each trial are summarized in Table S1.

Additionally, we generated a mapping population of 160 individuals by crossing the chloride-excluding accession *V. acerifolia* longii 9018 with the commercial rootstock GRN3. This population, designated 18113, was planted in the field, and a chloride exclusion screening—similar to the one conducted with the GWAS population—was performed.

### Leaf chloride content phenotyping

After 21 days of salt treatment, all leaves and petioles from each plant were harvested and stored in a paper bag. The samples were air-dried in a drying room at 50°C for 2 weeks, then ground into fine powder. Subsequently, 0.25 g of the fine powder was mixed with 25 mL of deionized water in screw-cap bottles, following the protocol of Heinitz et al. (2020). The mixture in screw-cap bottles was shaken at 150 revolutions per minute for 1 h, then filtered through an 11-µm filter to obtain a clear solution without leaf particles. The chloride content of the filtered samples was then measured using a silver ion titration chloridometer (Model 926, Nelson-Jameson Inc.) following the manufacturer’s calibration protocol with known chloride standards. Each sample was measured three times, and the average chloride concentration was expressed in mg/L.

The chloride content was analyzed using linear mixed models (LMMs) implemented in the *lme* package in R (Bates et al., 2015). We generated BLUPs for each accession using the following model:

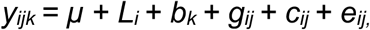

where *y_ijk_* represents the phenotype of response variable *i* in trial *j* and block *k*; *μ* is the overall mean; *L_i_* is the fixed effect of trial; *g_ij_* is the random genetic effect of regular individuals in trial *j*; *c_ij_* is the random genetic effect of experimental checks in trial *j*; and *e_ij_* is the random residual effect.

### Phylogenetic and population structure analysis

A phylogenetic analysis was performed to infer evolutionary relationships among accessions. A neighbor-joining tree was constructed based on genetic distance matrices allowing visualization of genetic divergence and the hierarchical relationships between the accessions using the R packages “*ape*” (Paradis et al., 2004) and “*ggtree*” (Yu et al., 2017). Population structure analysis was conducted using the sparse non-negative matrix factorization (sNMF) algorithm implemented in the “*LEA*” package in R (Frichot & François, 2015). The optimal number of genetic clusters (K) was determined by evaluating cross-entropy values. The analysis was finalized with K = 9, and the admixture proportions (Q-matrix) were extracted based on the results.

### Genotyping and GWAS

Sequencing data for the accessions used was compiled from previous work (Morales-Cruz et al., 2021; Morales-Cruz et al., 2023; Cochetel et al., 2023), and is available at NCBI under BioProject ID: PRJNA731597, PRJNA842753, and PRJNA984685. We filtered and processed raw sequencing reads using Trimmomatic-0.36 and FastQC. Reads were scanned in sliding windows of 4 base pairs, and bases were trimmed when the average quality per base dropped below Q20. We removed leading and trailing bases with quality scores below Q3 and retained only reads that were ≥60 bp after trimming. Filtered reads were then mapped to haplotype 1 of *V. girdiana* SC2 using the BWA-MEM algorithm implemented in bwa-0.7.8. Joint SNP calling was performed using GATK v.4.2.2.0. We first removed duplicate reads using the “MarkDuplicates” function from Picard tools, followed by the “AddOrReplaceReadGroups” function to label reads by individual. For SNP prediction, we applied the HaplotypeCaller algorithm with a ploidy of 2 and a mapping base quality score threshold of Q20. The final SNP calls were generated by combining VCF files from all individuals using the “GenotypeGVCFs” function with default parameters. SNP filtering was conducted using bcftools v1.9 and vcftools v0.1.17, applying stringent criteria to retain high-confidence variants. We kept only biallelic SNPs with, quality score >30, depth of coverage >5 reads, no more than 3× the median coverage depth across accessions, up to 25% missing data among individuals, and minor allele frequency > 0.1.

Genome-wide association was conducted using a mixed linear model (MLM) implemented in the gemma v.0.98.3 (Zhou & Stephens, 2012). The genomic relationship matrix (K) and 10 PCs, derived from SNP marker genotypes, were used to adjust for genetic relationships among individual. Prior to GWAS, PLINK v1.9 (Chang et al., 2015) was used for SNP pruning to reduce LD among markers using “--indep-pairwise 50 5 0.2”. The pruned dataset was used to generate the standardized relatedness matrix to account for kinship among individuals. The significance threshold was established using false discovery rate (FDR) correction to identify SNPs that exceeded the genome-wide significance level. To pinpoint potential candidate genes within each QTL region, we leveraged positional gene information and functional annotations from our dataset (www.grapegenomics.com). Candidate genes were identified within a ±100 kb window surrounding each QTL.

### QTL mapping in the longii 9018 **×** GRN population

Genomic DNA was extracted from young leaf samples at the University of Minnesota Genome Center, while genotyping-by-sequencing (GBS) was performed at the University of California, Davis Genome Center using the Illumina NovaSeq 6000 platform with paired-end (PE) 150 bp reads and an average sequencing depth of 10× coverage. Variant calling was conducted using TASSEL (Glaubitz et al., 2014), with haplotype 1 of *V. acerifolia* longii 9018 as the reference genome. To ensure high-quality variant calls, reads with a depth below 20 or above 200 were marked as missing, along with heterozygous calls exhibiting skewed allele proportions. Markers with >25% missing data and individuals with >40% missing genotype information were excluded from further analysis. Additionally, distorted markers were removed based on a chi-square test at p < 0.05.

Two genetic maps, one for longii 9018 and another for GRN3, were constructed using a pseudo-test cross linkage mapping strategy (Grattapaglia and Sederoff, 1994) in BatchMap (Schiffthaler et al., 2017). To generate predictors for the progeny population, we applied a similar model to the one used for GWAS BLUP estimation. A single-marker regression approach was implemented in R/qtl (Broman et al., 2003), with genome-wide significance thresholds established through 1000 permutation tests.

## ACKNOWLEDGMENTS

The authors would like to thank Sabrina Colacion, Mikayla Bailey, and Guillermo Garcia-Zamora for their support with greenhouse experiments and vineyard maintenance.

## DATA AVAILABILITY STATEMENT

All the collected data have been made available in the supplementary files accompanying this manuscript.

---

**Figure S1.**
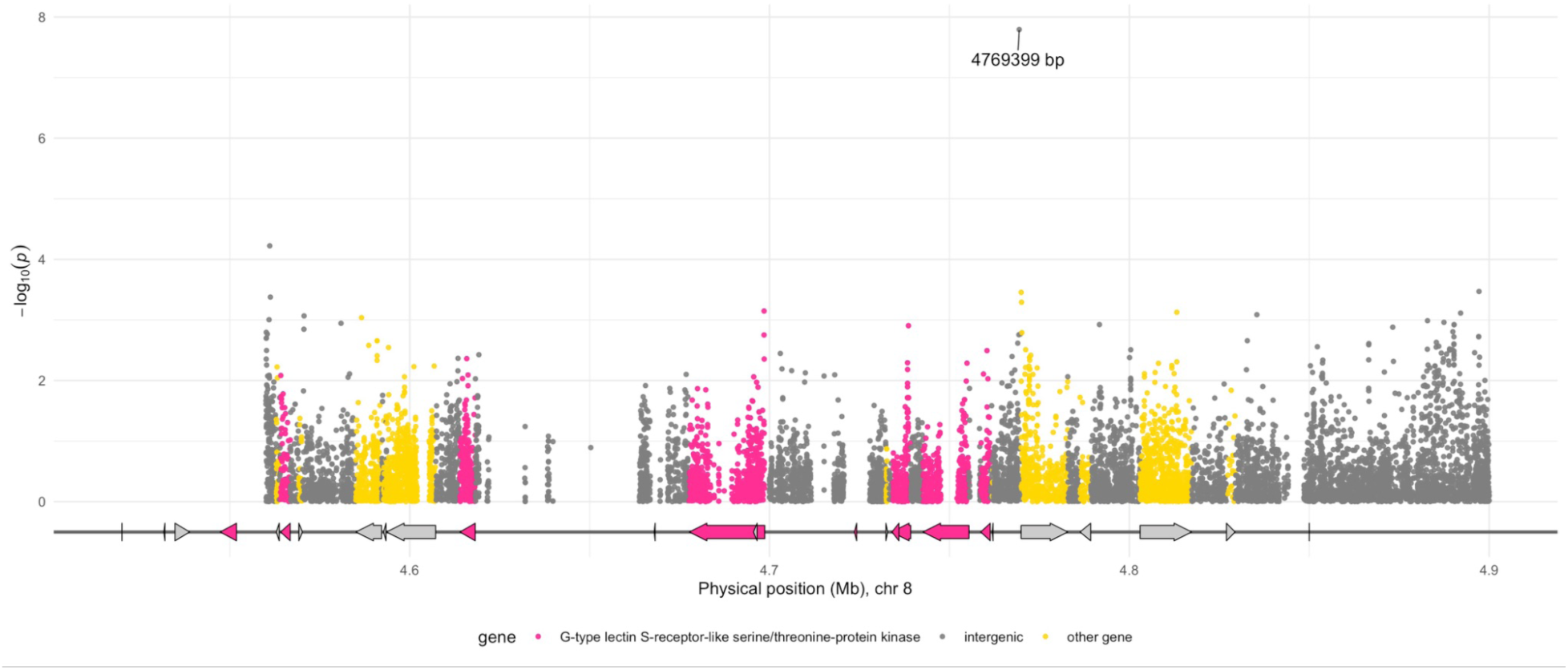
Zoomed view of the QTL in chromosome 19 (4.56-4.9 Mb).

